# Development and Application of a Filamentous Phage-Based Rapid Detection Tool for *Ralstonia pseudosolanacearum*

**DOI:** 10.1101/2025.08.15.670626

**Authors:** Qiaoxian Luo, Fangling Shu, Dehong Zheng

## Abstract

Bacterial wilt caused by the *Ralstonia solanacearum* species complex (RSSC) is a devastating plant disease with a broad host range. Early detection is critical for disease management, yet conventional methods lack speed and specificity. This study developed a rapid detection system using engineered filamentous phages (RSCq) expressing bioluminescence genes (*luxAB, nanoluc*, and *luxSit-i*). Among the constructs, RSCqluxAB demonstrated optimal performance, detecting *R. pseudosolanacearum* at 1.58×10^3^ CFU/mL within 12 hours, with minimal background noise. The phage retained stability for six months at 4°C, proving suitable for field applications. These findings highlight its potential for early pathogen monitoring, quarantine enforcement, and precision agriculture, though further validation in soil/plant samples is needed.

## Introduction

Bacterial wilt, caused by *Ralstonia solanacearum* species complex (RSSC), is a devastating soil-borne disease and one of the most severe bacterial plant diseases, often referred to as the “cancer” of plants (Mansfield *et al*., 2012). RSSC, comprising *R. pseudosolanacearum, R. solanacearum*, and *R. syzygii* (Etminani et al., 2020), has an extremely broad host range, infecting over 400 plant species across 50 families, with Solanaceae plants being the most severely affected (Lowe-Power *et al*., 2024). It also damages important woody plants such as olive, eucalyptus, and mulberry (Yuan *et al*., 2023). Additionally, RSSC is widely distributed geographically (EPPO, 2025), highly resilient, and can survive for years in moist environments (Alvarez *et al*., 2008). Its remarkable genetic diversity and propensity for mutation further complicate disease control, making bacterial wilt extremely difficult to manage, with limited effective control measures currently available (Yang *et al*., 2023).

In the early stages of infection, plants show no obvious symptoms. However, once the disease erupts, plants rapidly wilt and die within a short period. Chemical interventions post-outbreak have limited efficacy, making early detection of the pathogen during the initial planting phase critical for precise disease management (Fan *et al*., 2023). For example, screening potato seed tubers for RSSC can prevent the use of infected seeds, reducing the incidence of bacterial wilt at its source (Okiro *et al*., 2019). Beyond parasitizing host plants, RSSC can weakly colonize non-host plants and survive saprophytically in soil. Detecting the pathogen in soil or non-host plants before planting susceptible crops can effectively predict disease outbreaks.

Moreover, detecting RSSC is essential for distinguishing it from diseases with similar symptoms. For instance, tobacco is threatened by four major soil-borne diseases: bacterial wilt (caused by RSSC), blackleg disease (caused by *Pectobacterium carotovorum*) (Xia & Mo, 2007), black shank (caused by *Phytophthora nicotianae*) (Liu *et al*., 2025), and root rot (caused by *Fusarium oxysporum*) (Gangwar *et al*., 2025). These diseases exhibit similar wilting symptoms, often leading to misdiagnosis. Since the pathogens belong to different taxonomic groups (bacteria, fungi, and oomycetes), their control requires distinct chemical treatments. Incorrect diagnosis results in ineffective pesticide use and failed control measures. Thus, accurate and rapid diagnosis of tobacco bacterial wilt is crucial for effective management.

Due to its significant threat to agriculture, RSSC has been listed as an A2-regulated pest by the European and Mediterranean Plant Protection Organization (EPPO) and other countries, including the UK and Brazil (EPPO, 2025). Notably, banana bacterial wilt (caused by *R. solanacearum* race 2) is also listed as a quarantine pest (No. 324) in China’s List of Quarantine Pests for Imported Plants (2021). Therefore, rapid, simple, and effective detection methods for RSSC can enhance quarantine capabilities, reduce misdiagnoses, improve customs efficiency, and safeguard agricultural production.

## Results

### 1. Construction of Reporter Phages and Evaluation of Detection Performance for *R. pseudosolanacearum*

RSCqluxA and RSCqluxB are engineered filamentous phages based on RSCq, expressing the *luxA* and *luxB* genes, respectively (Peng *et al*., 2024) (Figure 1A). LuxA and LuxB form a heterodimer that catalyzes a luminescent reaction in the presence of long-chain aldehyde substrates (Lin *et al*., 1993). Our previous studies demonstrated that *R. pseudosolanacearum* co-infected with RSCqluxA and RSCqluxB and supplemented with decanal emits detectable bioluminescence. This study aimed to develop a rapid detection system for RSSC based on this property.

**Figure 1.**
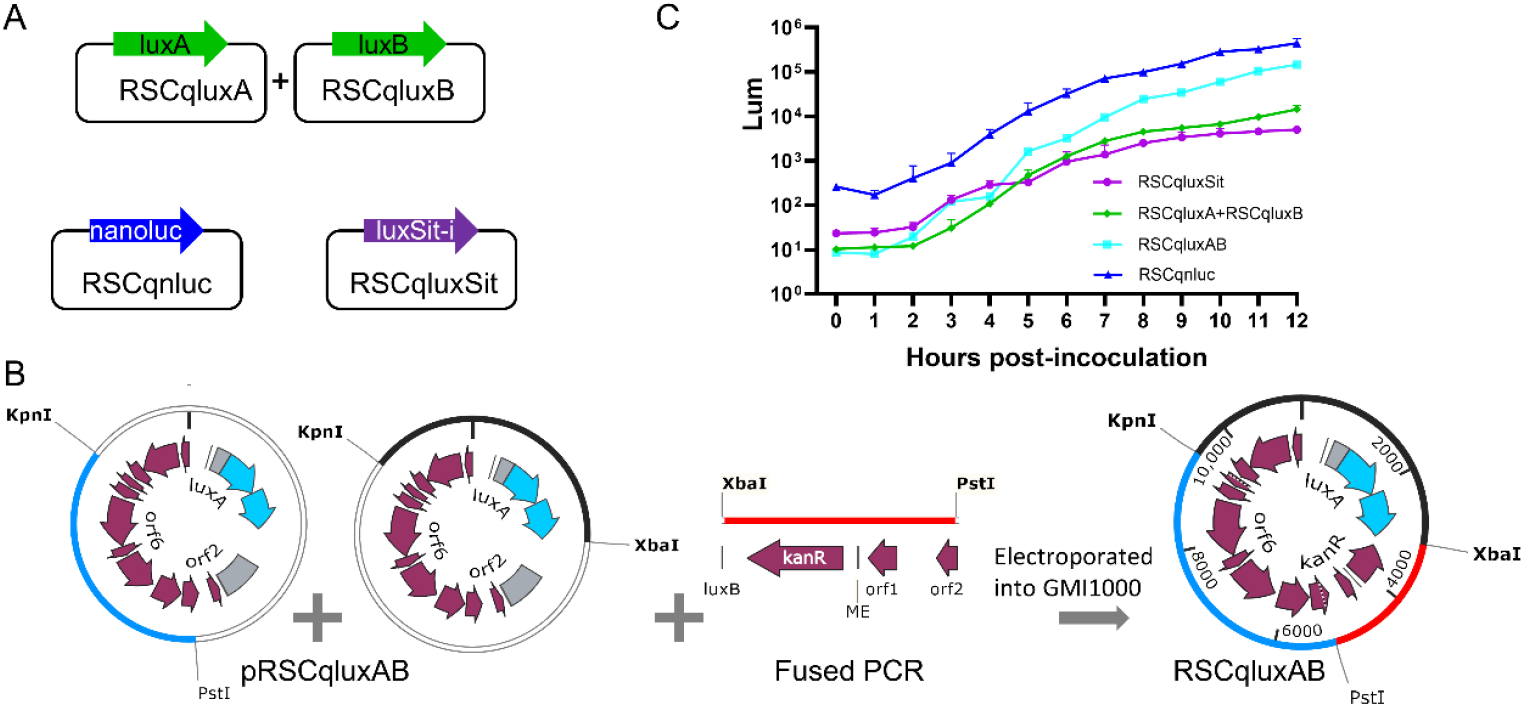
Reporter phages construction and *R. pseudosolanacearum* detection evaluation. **A**. Schematics of reporter phage RSCqluxA+RSCqluxB, RSCqnluc, and RSCqluxSit. **B**. Construction of reporter phage RSCqluxAB. **C**. Luminescence kinetics of reporter phages -infected *R. pseudosolanacearum*.

To compare the performance of different luminescence systems, we replaced the *luxA* gene in the engineered phages with the small, high-performance nanoluc luciferase gene (England *et al*., 2016) and the luxSit-i gene designed via generative AI (Yeh *et al*., 2023), yielding reporter phages RSCqnluc and RSCqluxSit, respectively (Figure 1A). To improve the LuxAB system’s efficiency, we cloned both luxA and luxB into RSCq. However, transforming *R. pseudosolanacearum* with the plasmid pRSCqluxAB did not yield functional phages, likely due to the large insert size (luxA + luxB, 2.1 kb). By removing the *E. coli* replication origin (R6Kγori) from pRSCqluxAB via multi-fragment ligation, we obtained the infectious reporter phage RSCqluxAB (Figure 1B). We then infected *R. pseudosolanacearum* GMI1000 (initial concentration: 10^7^ CFU/mL) with RSCqluxA + RSCqluxB, RSCqnluc, RSCqluxSit, or RSCqluxAB and measured luminescence at different time points. As shown in Figure 1C, RSCqnluc-infected *R. pseudosolanacearum* exhibited the strongest luminescence after 12 hours, followed by RSCqluxAB-infected *R. pseudosolanacearum*. However, RSCqnluc also showed high background luminescence at 0 hours, and its substrate (Fluorofurimazine) is expensive. Thus, we selected RSCqluxAB for further study.

### 2. Detection Limit and Storage Stability of RSCqluxAB

We assessed the *R. pseudosolanacearum* detection sensitivity of phage RSCqluxAB by incubating it with *R. pseudosolanacearum* GMI1000 at varying concentrations and monitoring luminescence. As shown in Figure 2A, the system exhibited minimal background luminescence in the absence of *R. pseudosolanacearum*. Higher bacterial concentrations yielded faster luminescence detection. RSCqluxAB detected 1.58 × 10^7^ CFU/mL after 3 hours and 1.58 × 10^3^ CFU/mL after 12 hours.

**Figure 2.**
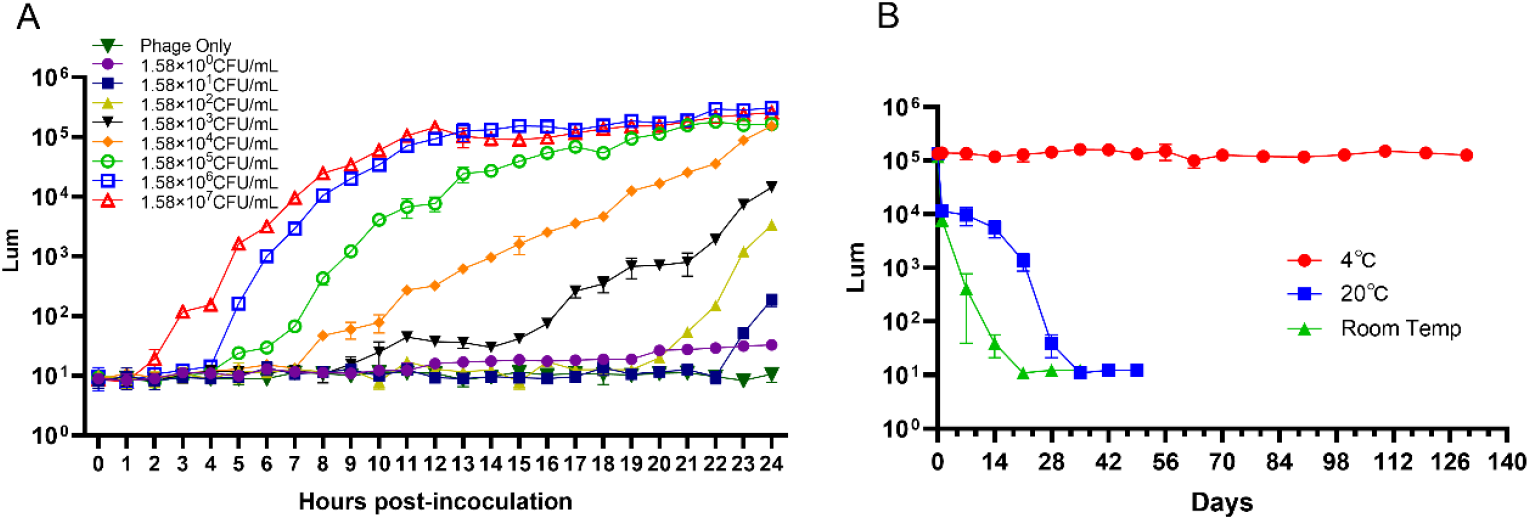
Detection limit and storage stability of RSCqluxAB. **A.** Luminescence at varying RSSC concentrations and time points. **B**. Luminescence after 12-hour incubation with 10^7^ CFU/mL RSSC following storage at different temperatures.

Storage stability is critical for pathogen detection systems. We stored the filamentous phage RSCqluxAB at three different temperature conditions (4°C refrigerator, 20°C incubator, and room temperature) and periodically evaluated its detection efficacy against *R. pseudosolanacearum*. RSCqluxAB stored at 4°C maintained stable activity for six months, while phages stored at 20°C or room temperature lost detection capability within a month (Figure 2B).

## Materials and Methods

### 1. Strains and Culture Conditions

*R. pseudosolanacearum* strains were cultured in BG medium 10 g/L (bactopeptone, 1 g/L casamino acids, 1 g/L yeast extract, and 5 g/L glucose) at 28°C (Perrier *et al*., 2018). *Escherichia coli* DH5αλpir was used for engineered filamentous phage plasmid construction, with kanamycin (25 μg/mL) added as needed.

### 2.Reporter Phage Construction

RSCqluxA and RSCqluxB were constructed previously (Peng et al., 2024). The nanoluc and luxSit-i genes replaced *luxA* in RSCqluxA to generate pRSCqnluc and pRSCqluxSit, which were electroporated into *R. pseudosolanacearum* GMI1000 to yield RSCqnluc and RSCqluxSit. The *luxAB* operon replaced *luxA* in RSCqluxA to generate recombinant plasmid pRSCqluxAB. The plasmid pRSCqluxAB was digested with *Kpn*I and *Pst*I to obtain a 4.3 kb DNA fragment, and with *Kpn*I and *Xba*I to yield a 4.9 kb DNA fragment. A 2 kb DNA fragment lacking the *E. coli* replication element was amplified by overlap PCR using pRSCqluxAB as template. These three fragments were ligated and directly electroporated into *R. pseudosolanacearum* GMI1000. The reporter filamentous phage RSCqluxAB was subsequently isolated from the culture supernatant of transformants.

### 3. Bioluminescence Detection

The *R. pseudosolanacearum* strains harboring filamentous phages were cultured, and the corresponding reporter filamentous phages were harvested from the culture supernatant. A 1% (v/v) aliquot of the reporter phages was added to BG medium containing *R. pseudosolanacearum* at varying concentrations. After incubation for specified durations, the corresponding detection substrates were added, and bioluminescence values were quantified using a microplate reader. Substrates: decanal (for RSCqluxA+RSCqluxB and RSCqluxAB), Furimazine (for RSCqnluc), and DTZ (for RSCqluxSit).

## Discussion

Compared to traditional methods, phage-based detection is simpler, cost-effective, and distinguishes live/dead bacteria (Meile *et al*., 2023, Braun *et al*., 2023). RSCqluxAB detected 10^7^ CFU/mL in 3 hours and 10^3^ CFU/mL in 12 hours, showing promise for rapid detection. Challenges remain, including improving sensitivity, reducing reliance on equipment, and validating performance in soil/plant samples.

